# *Enterococcus faecalis* strains with compromised CRISPR-Cas defense emerge under antibiotic selection for a CRISPR-targeted plasmid

**DOI:** 10.1101/220467

**Authors:** Wenwen Huo, Valerie J. Price, Ardalan Sharifi, Michael Q. Zhang, Kelli L. Palmer

## Abstract

*Enterococcus faecalis* is a Gram-positive bacterium that natively colonizes the human gastrointestinal tract and opportunistically causes life-threatening infections. Multidrug-resistant (MDR) *E. faecalis* strains have emerged that are replete with mobile genetic elements (MGEs). Non-MDR *E. faecalis* strains frequently possess CRISPR-Cas systems, which reduce the frequency of mobile genetic element (MGE) acquisition. We demonstrated in previous studies that *E. faecalis* populations can transiently maintain both a functional CRISPR-Cas system and a CRISPR-Cas target. In this study, we used serial passage and deep sequencing to analyze these populations. In the presence of antibiotic selection for the plasmid, mutants with compromised CRISPR-Cas defense and enhanced ability to acquire a second antibiotic resistance plasmid emerged. Conversely, in the absence of selection, the plasmid was lost from wild-type *E. faecalis* populations, but not *E. faecalis* populations that lacked the *cas9* gene. Our results indicate that *E. faecalis* CRISPR-Cas can become compromised under antibiotic selection, generating populations with enhanced abilities to undergo horizontal gene transfer.

**Importance:** *Enterococcus faecalis* is a leading cause of hospital-acquired infections and disseminator of antibiotic resistance plasmids among Gram-positive bacteria. We have previously shown that *E. faecalis* strains with an active CRISPR-Cas system can prevent plasmid acquisition and thus limit the transmission of antibiotic resistance determinants. Yet, CRISPR-Cas was not a perfect barrier. In this study, we observed populations of *E. faecalis* with transient co-existence of CRISPR-Cas and one of its plasmid targets. Our experimental data demonstrate that antibiotic selection results in compromised *E. faecalis* CRISPR-Cas function, thereby facilitating the acquisition of additional resistance plasmids by *E. faecalis*.

## Introduction

*Enterococcus faecalis* is a Gram-positive bacterium that natively colonizes the human gastrointestinal tract and opportunistically causes life-threatening infections (1–4). Antibiotic resistance is a major concern for treatment of these infections. *E. faecalis* can acquire antibiotic resistance through horizontal gene transfer (HGT), mediated primarily by plasmids, transposons, and integrative conjugative elements (5–7). One of the most clinically relevant forms of HGT in *E. faecalis* is afforded through the pheromone-responsive plasmids. The pheromone-responsive plasmids are narrow host range conjugative plasmids that disseminate antibiotic resistance and virulence genes among *E. faecalis* (5, 8–10) and mobilize resistance genes to other pathogens (2, 11–14).

Some *E. faecalis* encode CRISPR-Cas systems. We have demonstrated that these systems reduce the spread of antibiotic resistance plasmids among *E. faecalis in vitro,* and *in vivo* in the murine intestine (15–19). CRISPR-Cas systems confer sequence-specific genome defense against mobile genetic elements (MGEs) (20, 21). Type II CRISPR-Cas systems occur in *E. faecalis* (22–24). They consist of a CRISPR array and CRISPR-associated (*cas*) genes. In *E. faecalis*, the CRISPR is an array of 36 base pair (bp) repeat sequences interspersed by 30 bp sequences referred to as spacers. When cells are challenged with a MGE, some incorporate a short segment (protospacer) of the invading MGE genome into the CRISPR as a novel spacer (20, 25). By this mechanism, the CRISPR serves as a heritable molecular memory of MGE encounters. Short sequence motifs adjacent to protospacers, called protospacer adjacent motifs (PAMs), as well as the Cas9, Cas1, and Cas2 nucleases are important for adaptation (26–31). The CRISPR is transcribed and processed into small RNAs called crRNAs; one crRNA corresponds to one spacer (32). If a MGE possessing the protospacer and PAM enters the cell, the Cas9 nuclease is directed to the MGE genome by the crRNA. Cas9 cleaves the invading MGE, generating a double-stranded DNA break, thereby inhibiting its entry (33–36).

Several of our studies (17, 18, 37) have focused on the CRISPR3-Cas locus of the *E. faecalis* strain T11RF, which natively lacks horizontally acquired antibiotic resistance genes (38, 39). *E. faecalis* T11RF is closely related to the model multidrug-resistant *E. faecalis* strain V583, but lacks the ∼620 kb of horizontally acquired genome content that V583 possesses (39). T11RF has a CRISPR-Cas system, while V583 does not (24). In general, multidrug-resistant *E. faecalis* lack CRISPR-Cas. The *E. faecalis* T11RF CRISPR3-Cas possesses 21 spacers. Spacer 6 is identical to the *repB* gene of the model pheromone-responsive plasmid pAD1. pAD1 encodes a haemolytic bacteriocin referred to as cytolysin and the virulence factor aggregation substance (8, 40). We previously demonstrated that T11RF CRISPR3-Cas interferes with acquisition of the pAD1 derivative pAM714, which encodes erythromycin resistance via *ermB* on Tn*917* (41, 42). Deletion of *cas9* or spacer 6 in *E. faecalis* T11RF resulted in up to a 150-fold increase in pAM714 acquisition in *in vitro* mating with *E. faecalis* OG1SSp(pAM714) donors (17). This increase in conjugation frequency was not observed for the pheromone-responsive plasmid pCF10, which is not natively targeted by the T11RF CRISPR3 spacers (17). These experiments confirmed that CRISPR3-Cas is a sequence-specific anti-plasmid defense system in *E. faecalis* T11RF.

Despite *E. faecalis* T11RF CRISPR3-Cas acting as a barrier to pAM714, up to 10^5^ T11RF(pAM714) transconjugants still arose in *in vitro* conjugation reactions (17). In this study, we tested the investigated the fate of these transconjugants in serial passage experiments with and without selection for the plasmid.

## Results

### Serial passage experiments to study transconjugant evolution over time

We mated *E. faecalis* OG1SSp(pAM714) donors and T11RF or T11RF Δ*cas9* recipients for 18 hours on agar plates without erythromycin selection (17). After 18 hours, mating reactions were recovered and plated on agar selective for transconjugants, donors, and recipients. These data were previously reported and demonstrated that CRISPR-Cas significantly reduces pAM714 acquisition by T11RF (17). We randomly selected 6 T11RF(pAM714) and 6 T11RFΔ*cas9*(pAM714) transconjugants for further analysis in this study. The 6 T11RF(pAM714) transconjugants are referred to hereafter as WT1-WT6, and the 6 T11RFΔ*cas9*(pAM714) transconjugants are referred to as Δ1-Δ6. The transconjugant colonies were each resuspended in brain heart infusion (BHI) broth. Samples were removed from the colony resuspensions for serial dilution and plating to determine the percentage of erythromycin-resistant cells, and for PCR and sequencing to determine the size and sequence of the CRISPR3. These data are referred to as “Day 0.” Next, each transconjugant colony resuspension was split equally into BHI broth and BHI broth with erythromycin and passaged daily for 14 days. At every passage, broth samples were removed to determine the percentage of erythromycin-resistant cells and the size (by agarose gel electrophoresis) and sequence (by Illumina sequencing) of the CRISPR3 amplicon. As a control, wild-type T11RF was also passaged for 14 days in BHI broth.

At Day 0, the WT1-WT6 populations, except for WT4, were a mix of erythromycin-sensitive (primarily) and erythromycin-resistant cells (Figure S1A). The Δ1-Δ6 populations were comprised of erythromycin-resistant cells (Figure S1A). The CRISPR3 arrays of all populations were wild-type size (1.76 kb) based on PCR amplification and agarose gel electrophoresis analysis of products (Figure S1B).

### Erythromycin resistance is lost from most WT but no Δ*cas9* populations during passage in non-selective medium

The frequency of erythromycin-resistant cells decreased in the WT1, WT2, WT3, WT5, and WT6 populations over the course of serial passage in BHI broth lacking erythromycin (Figure 1A). In contrast, erythromycin resistance was stably maintained at high frequencies in the WT4 population and all of the Δ1-Δ6 populations in the absence of erythromycin selection (Figure 1A). The CRISPR3 arrays of all populations were of wild-type size based on PCR amplification and agarose gel electrophoresis analysis of products (Figure 1B and Figure S2).

**Figure 1.**
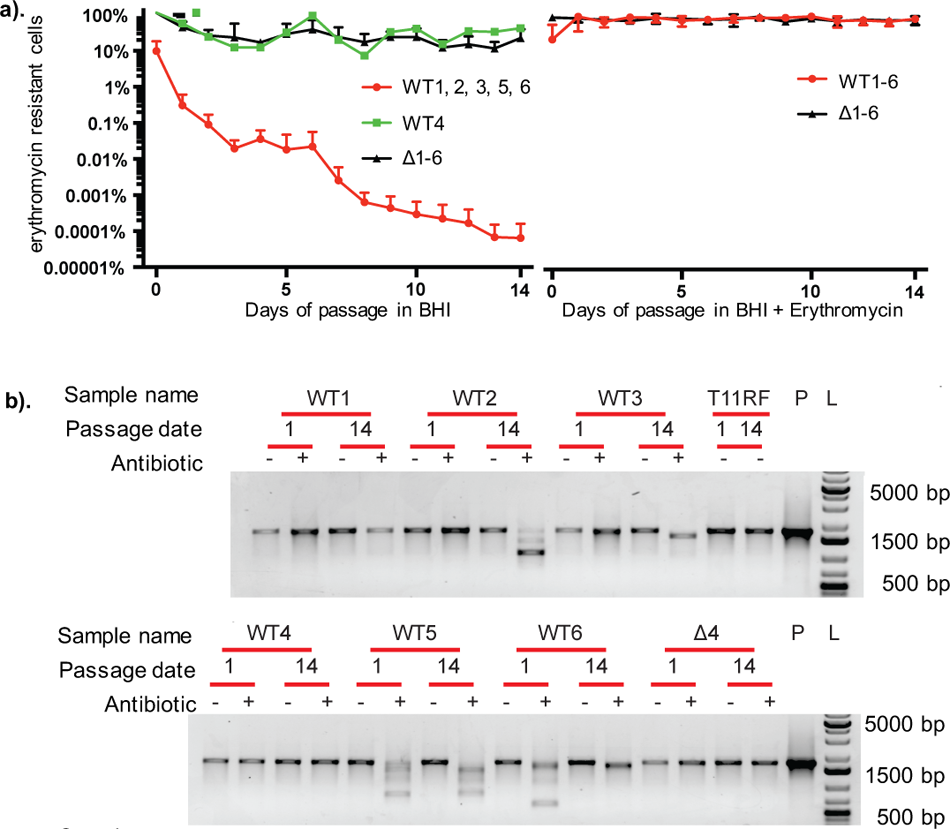
Antibiotic resistance phenotypes and qualitative assessment of CRISPR3 genotypes in serially passaged transconjugant populations. a) Erythromycin resistant CFU/mL (expressed as percent of total CFU/mL) remaining in *E. faecalis* populations over the course of passage without (left) and with (right) antibiotic selection. WT transconjugant populations are shown in green or red and τι*cas9* transconjugant populations are shown in black. b) CRISPR3 amplicon size from early (Day 1) and late (Day 14) passage dates for six WT transconjugant populations and a representative Δ*cas9* transconjugant population (Δ4). The CRISPR3 region was PCR-amplified using aliquots from passaged cultures as template. As a control, T11RF without pAM714 was passaged for 14 days and the CRISPR3 locus amplified (T11RF). P: positive control, T11RF genomic DNA. L: DNA ladder.

We analyzed the WT4 population further because it was an outlier compared to the other WT populations. We sequenced the *cas9* coding region of the WT4 population from Day 0 and identified a mutation resulting in an Ala749Thr substitution. Ala749 occurs within the RuvC nuclease domain and is conserved in the well-studied Cas9 of *Streptococcus pyogenes* (17). The Ala749Thr substitution may result in loss in Cas9 function, causing WT4 to phenocopy the Δ1-Δ6 populations.

### CRISPR-Cas mutants emerge in WT populations during passage with erythromycin

We observed stable maintenance of erythromycin resistance in all WT1-WT6 and Δ1-Δ6 populations during passage with erythromycin selection (Figure 1A). By Day 14 of passage with erythromycin, CRISPR3 arrays in all WT populations except WT1 and WT4 were smaller in size than the wild-type T11RF control (Figure 1B). The variation in array size initiated sporadically over the 14 days, and each transconjugant had a unique pattern (Figure S2). Sanger sequencing of the CRISPR3 array amplicons from Day 1 and Day 14 erythromycin-passaged populations revealed that the CRISPR3 spacer 6 was either deleted from the array or the spacer 6 region had poor sequence quality (Table 2). We will refer to spacer 6 as S_6_ hereafter, and use this subscript nomenclature for the other 20 spacers of the T11RF CRISPR3 array (e.g., S_1_, S_2_, etc.).

**Table 1.** Bacterial strains and plasmids used.

| Name | Description | Reference |
| --- | --- | --- |
| ***E. faecalis* strains** |  |  |
| T11RF | Rifampicin- and fusidic acid-resistant derivative of the human urine isolate T11 | (17, 39) |
| T11RFΔcas9 | Derivative of T11RF with *cas9* deleted | (17) |
| OG1RF | Rifampicin- and fusidic acid-resistant derivative of the human oral isolate OG1 | (22, 67) |
| OG1SSp | Spectinomycin- and streptomycin-resistant derivative of OG1; donor strain for conjugation assays | (41-43) |
| **Plasmids** |  |  |
| pAM714 | 65 kb PRP encoding erythromycin on Tn917, derivative of pAD1 | (41) |
| pCF10 | 67 kb PRP encoding tetracycline resistance on Tn925 | (43) |
| pLZ12 | Broad host range shuttle vector encoding chloramphenicol resistance | (68) |
| pKH12 | pLZ12 with *oriT* | (16) |
| pKHS5 | pKH12 with CRISPR2 protospacer S5 and CRISPR1/2 PAM | This study |
| pKHS96 | pKH12 with CRISPR1 protospacer S96 and CRISPR1/2 PAM | (16) |
| pVP107 | pLT06 with T11CR2 protospacer and CRISPR1/2 PAM | (17) |
| pWH107 | pVP107 digested with XbaI/SphI and re-ligated with primers pVP107_XbaI_For/pVP107_SphI_Rev to remove PstI enzyme site | This study |
| pWH107.S1 | pWH107 with T11CR3 protospacer S1 and CRISPR3 PAM inserted between BamHI/PstI | This study |
| pWH107.S6 | pWH107 with T11CR3 protospacer S6 and CRISPR3 PAM inserted between BamHI/PstI | This study |
| pWH107.S7 | pWH107 with T11CR3 protospacer S10 and CRISPR3 PAM inserted between BamHI/PstI | This study |

**Table 2.** CRISPR alleles.

| Sample name | Day 1 Sanger<sup>c</sup> | Day 14 Sanger<sup>c</sup> | Day 1 Amplicon<sup>d</sup> | Day 14 Amplicon<sup>d</sup> |
| --- | --- | --- | --- | --- |
| T11RF control<sup>a</sup> | WT | WT | WT | WT |
| $\Delta 4^b$ | WT | WT | WT | WT |
| WT1<sup>b</sup> | Poor quality at S<sub>6</sub>-<br>S<sub>7</sub> | WT | WT | WT |
| WT2<sup>b</sup> | WT | $\Delta S_2-S_{11}$ | WT | $\Delta S_2-S_{11}$ |
| WT3<sup>b</sup> | Poor quality at S<sub>6</sub>-<br>S<sub>7</sub> | $\Delta S_5-S_7$ | WT | $\Delta S_5-S_7$ |
| WT5<sup>b</sup> | Poor quality at S<sub>3</sub>-<br>S<sub>8</sub> | Poor quality at S<sub>3</sub>-<br>S<sub>9</sub> | WT | $\Delta S_5-S_8$ |
| WT6<sup>b</sup> | Poor quality at S<sub>1</sub>-<br>S<sub>7</sub> | Poor quality at S<sub>5</sub>-<br>S<sub>6</sub> | $\Delta S_5-S_6$ | $\Delta S_6$ |

<sup>a</sup>T11RF without pAM714 passaged for 14 days in BHI medium.
<sup>b</sup>pAM714 transconjugants passaged for 14 days in BHI medium with erythromycin.
<sup>c</sup>CRISPR3 alleles detected by Sanger sequencing.
<sup>d</sup>CRISPR3 alleles detected by Illumina amplicon deep sequencing. Abundant alleles with p-value < 0.05 are shown.

In contrast to WT populations, CRISPR3 array sizes for the Δ1-Δ6 populations were unchanged after passage in medium with erythromycin (Figure 1B and Table 2). We chose the Δ4 population as a representative of the Δ1-Δ6 populations for future analyses.

To investigate whether mutations were present outside the CRISPR3 array, we performed whole genome Illumina sequencing on select populations. We observed variation in the *cas9* sequence of the WT1, WT2, and WT3 Day 14 populations relative to the T11RF wild-type (Table 3). All of the mutations led to nonsynonymous changes (Table 3). Some of the sequenced populations possessed variations in one or more additional genes (Table S2). No mutations were identified in the S_6_ protospacer or the PAM region of the *repB* gene in pAM714. However, we did identify a mutation within *repB*, not associated with the protospacer or PAM, in the WT2 population (Table S2). We observed no evidence of Tn*917* movement from pAM714 into the T11 chromosome as all reads overlapping the ends of Tn*917* also overlapped the pAD1 reference sequence.

**Table 3.**
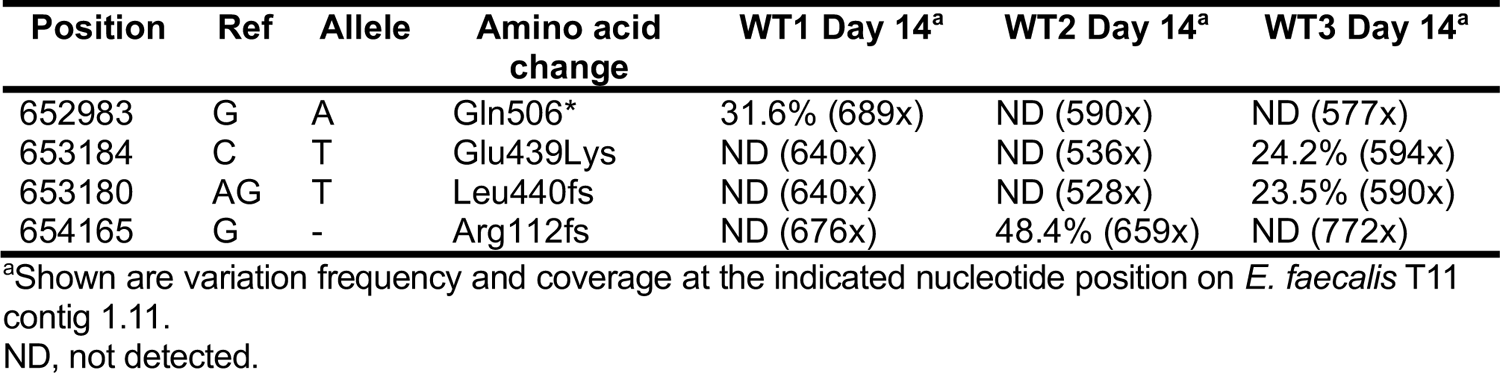
Nonsynonymous *cas9* mutations detected by whole genome sequencing.

| Position | Ref | Allele | Amino acid change | WT1 Day 14<sup>a</sup> | WT2 Day 14<sup>a</sup> | WT3 Day 14<sup>a</sup> |
| --- | --- | --- | --- | --- | --- | --- |
| 652983 | G | A | Gln506* | 31.6% (689x) | ND (590x) | ND (577x) |
| 653184 | C | T | Glu439Lys | ND (640x) | ND (536x) | 24.2% (594x) |
| 653180 | AG | T | Leu440fs | ND (640x) | ND (528x) | 23.5% (590x) |
| 654165 | G | - | Arg112fs | ND (676x) | 48.4% (659x) | ND (772x) |

<sup>a</sup>Shown are variation frequency and coverage at the indicated nucleotide position on *E. faecalis* T11 contig 1.11.
ND, not detected.

### CRISPR3 alleles with spacer deletions are present in most WT populations after serial passage with erythromycin selection

To attain greater resolution of CRISPR3 alleles, we used Illumina sequencing to analyze CRISPR3 amplicons. We analyzed the CRISPR3 amplicons obtained for the WT1-WT6 and Δ4 populations after Day 1 and Day 14 of serial passage with erythromycin. We also analyzed the wild-type T11RF control passaged in BHI broth. We achieved an average of 16 million reads for our amplicons (Table S3).

To identify specific CRISPR3 alleles in the amplicon sequencing, we created a pool of references by sliding a contiguous and non-overlapping window of 96 bp in length along CRISPR3 reference, which allows each reference to contain exactly two adjacent spacers connected by one repeat. The sliding window starts at 30 bp upstream of the first repeat and ends at 30 bp downstream of the terminal repeat. In total, 22 references were generated to represent wild type alleles. Then, we manually constructed artificial CRISPR3 reference sequences for every possible spacer deletion and rearrangement event with a length of 96 bp. In total, 462 references were constructed to represent all mutant alleles. We then mapped CRISPR amplicon reads to wild-type and mutant references (see Figure S3). The mapping was performed with stringency to allow unique mapping only. After mapping, the number of mapped reads per 96-bp reference was calculated and normalized by the total number of mapped reads per all references. To further evaluate the normalized reads per reference, we calculated z-score using mean and standard deviation from the T11RF Day 1 and Day 14 mapping result. The higher the z-score, the more abundant reads per reference. The significance of z-score was calculated using t-test with a degree of freedom of 483. By applying a p-value cutoff of 0.05, we identified the most abundant CRISPR3 alleles in each population (Figure 2 and Table 2). Of the five T11RF transconjugant populations analyzed, all other than WT1 possessed at least one significantly enriched mutant CRISPR3 allele after 14 days of passage with antibiotic selection, and each of those alleles lacked S_6_ (Table 2).

**Figure 2.**
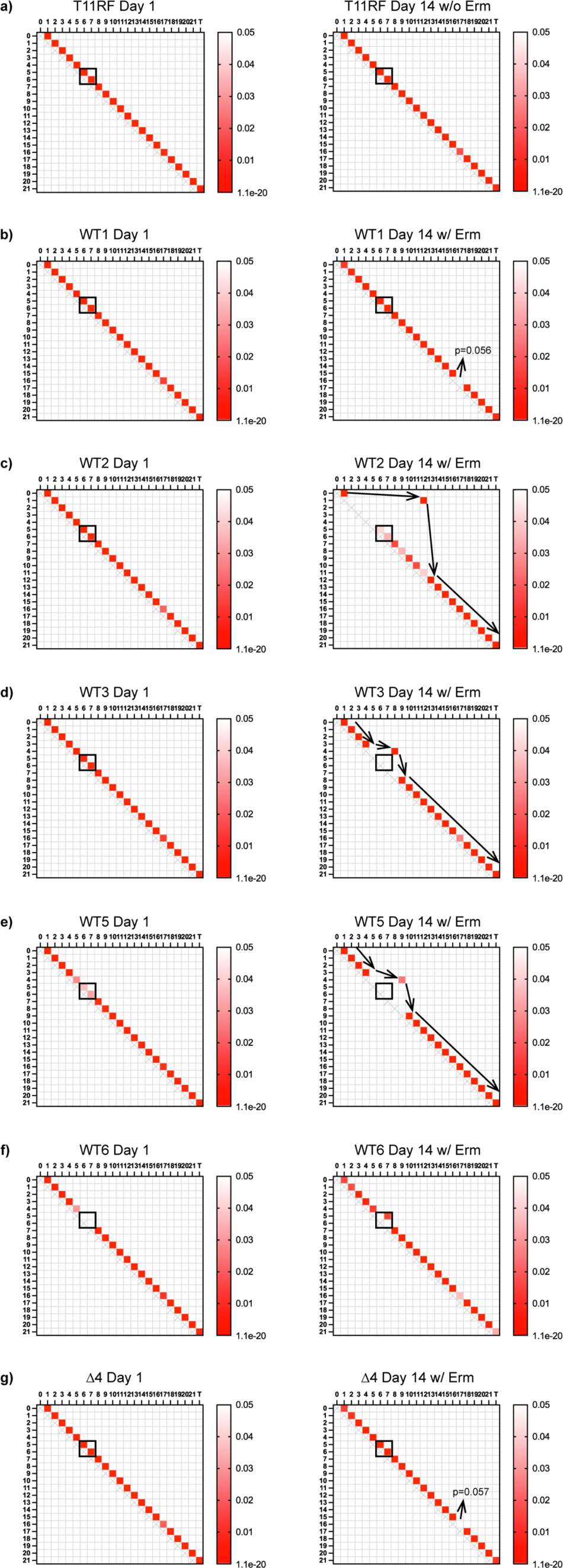
Detection of mutant CRISPR3 array alleles using amplicon sequencing. Heat maps show the mutant alleles with significant abundance (p-value <0.05) present at Day 1 and Day 14 of T11RF control (a) and antibiotic-passaged transconjugant populations (b-g). Each cell within heatmap represents one of the 484 artificial references where a 5’ spacer (rows) is linked with a 3’ spacer (columns) by a repeat. White cells represent mutant alleles that were not significantly abundant.

One drawback for our method is that it is difficult to set a p-value cutoff. Here we used canonical definition of 0.05. However, the normalized reads per reference for wild-type alleles is expected to be much higher than that from mutant alleles, and in fact, it is much higher in control samples (T11RF control Day 1 and Day 14). Such variation results in large standard deviation values which skew the calculation of z-scores and hence underestimates the abundance of mutant alleles. We predict that more mutant alleles existed in Day 14 WT transconjugant samples, which is supported by an examination of the normalized reads per reference plot (Figure S4), but we are uncertain about their significance as measured by p-value. Supporting our approach, control samples yielded results largely as expected. In control samples (T11RF and the representative Δ*cas9* transconjugant), only wild type alleles were significantly abundant with p-value < 0.05. However, we also observed uneven distribution among different 96-bp references representing wild-type alleles, which resulted in a “loss of spacer16-17” call for the Day 14 Δ*cas9* transconjugant at the p-value of 0.057. When looking at the normalized reads per reference heatmap (Figure S4), we believe spacer16-17 was still largely intact within the Δ*cas9* transconjugant population, the abundance of which is about 1000 times more than that from a mutant 96-bp reference. In short, our statistical approach has strengths and weaknesses, which have been discussed here.

### Functional assays confirm that CRISPR-Cas defense was compromised after antibiotic selection

Our sequencing analyses suggested that the WT transconjugants passaged with erythromycin had become deficient for CRISPR-Cas defense either by deletion of spacer 6 or inactivation of Cas9 by mutation. For a functional assessment, we tested whether the transconjugant populations could still defend against sequences targeted by T11RF CRISPR3-Cas. To do this, we utilized the pheromone-responsive plasmid pCF10 (43), which is not natively targeted by the T11RF CRISPR3-Cas system. We previously demonstrated that pCF10 transfer into wild-type and Δ*cas9* T11RF strains is equivalent (17). In this study, we modified pCF10 to be targeted by the T11RF CRISPR3-Cas system. We generated three derivatives of *E. faecalis* OG1SSp(pCF10), each with an insertion of a T11RF CRISPR3 spacer (S_1_, S_6_, or S_7_) and CRISPR3 consensus PAM in the pCF10 *uvrB* gene (Figure 3A). Disruption of *uvrB* does not impact pCF10 conjugation (17). We then compared conjugation of wild-type pCF10 and these derivatives into the control T11RF population that had been passaged in BHI broth for 14 days. As expected, conjugation frequencies were significantly lower for all pCF10 derivatives bearing CRISPR3 targets, as compared to the wild-type pCF10 (Figure 3B).

**Figure 3.**
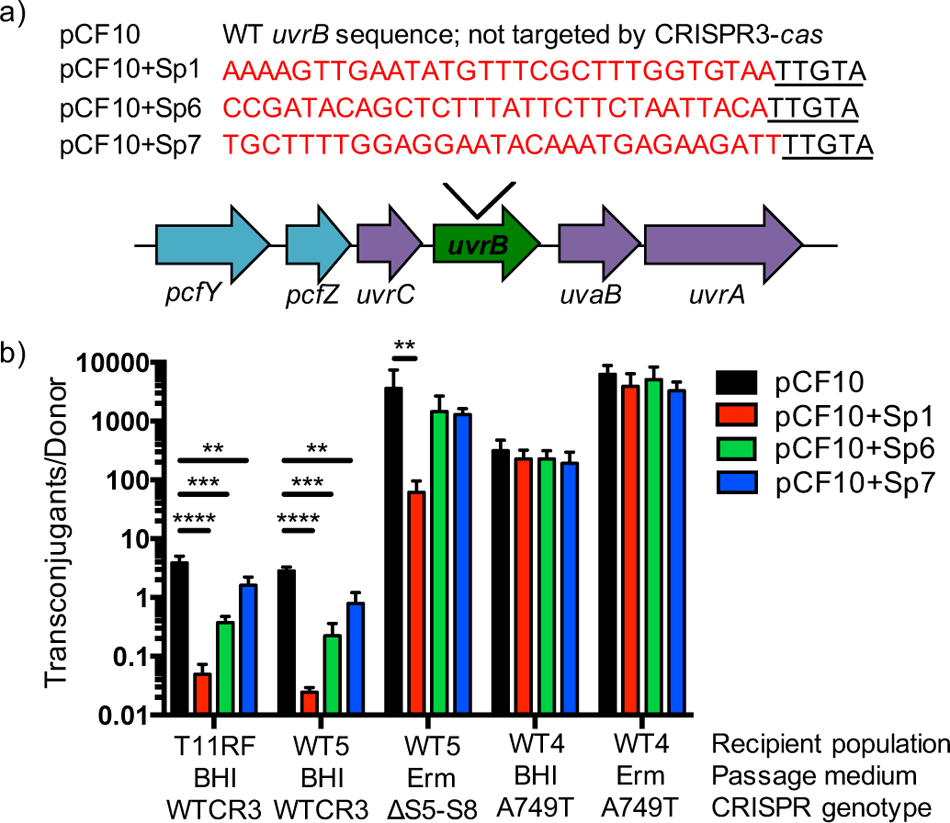
Experimental confirmation of compromised CRISPR-Cas function in antibiotic-passaged WT5 and in WT4. Day 14 transconjugant populations passaged in BHI and erythromycin (Erm) were used as recipients in conjugation reactions with OG1SSp(pCF10) and derivatives of pCF10. a) Schematic of where the T11 CRSPR3 spacer 1, spacer 6 and spacer 7 sequences with corresponding PAM (underlined) were introduced into pCF10. Wild-type pCF10 is not natively targeted by T11RF CRISPR3-Cas. b) Conjugation frequency (ratio of transconjugants to donors) from mating reactions. A749T refers to the *cas9* mutation detected in the WT4 population. Statistical significance was determined using a student’s t-test; P-values: ** ≤ 0.01; *** ≤ 0.001; **** ≤ 0.0001.

We then evaluated pCF10 transfer into the WT5 Day 14 populations. For WT5 passaged without erythromycin, we observed similar results to the T11RF control (Figure 3B), demonstrating that the CRISPR3-Cas system in this population is still functional. For WT5 passaged with erythromycin, conjugation frequency of only the pCF10 derivative bearing S_1_ was reduced relative to wild-type pCF10 (Figure 3B). This is consistent with the deletion of S_6_ and S_7_ in this population (Figure 2), and demonstrates that the sequencing data are accurate and not the result of PCR amplification bias. We note that we observed a ∼3 log higher conjugation frequency of pCF10 into this population compared to the WT5 population passaged without erythromycin (Figure 3B). This is due to comparatively low donor numbers (averages of 7.1 x 10^6^ and 5.6 x 10^9^ CFU/mL, respectively; raw transconjugant and donor numbers for Figure 2 are presented in Dataset S1). pAM714 encodes the bacteriocin, cytolysin (8). We previously reported that *E. faecalis* with pAM714 kill *E. faecalis* cells that lack pAM714 (18). The probable explanation is that the WT5 population passaged with erythromycin, which has a very high carriage rate of pAM714 (Figure 1) killed OG1SSp(pCF10) donors using cytolysin.

We also evaluated pCF10 transfer into the WT4 Day 14 populations. WT4 was unique among the 6 WT transconjugants in that it maintained erythromycin resistance at high frequencies in the absence of erythromycin selection (Figure 1A). We identified a probable loss-of-function mutation in the RuvC domain-encoding region of *cas9*. The WT4 populations did not interfere with any of the three pCF10 derivatives bearing CRISPR3 targets (Figure 3B). This is consistent with a loss of CRISPR-Cas function in the WT4 populations conferred by a loss of function mutation in *cas9*.

### Analysis of *E. faecalis* genomes identifies a strongly supported instance of *in situ* CRISPR-Cas compromise

Our data demonstrate that loss-of-function mutations in CRISPR-Cas arise that promote plasmid acquisition (for WT4) or plasmid maintenance (for WT2-3 and WT5-6) in *E. faecalis*. We used genome data available for *E. faecalis* to identify potential instances where this may have occurred in nature. We focused on published MDR *E. faecalis* strains that possess CRISPR-Cas, reasoning that CRISPR-Cas function may have been compromised, allowing for the accumulation of acquired resistance genes. More specifically, we narrowed our focus to strains that both encode CRISPR-Cas as well as resistance to the last-line antibiotics vancomycin or linezolid, which are often plasmid-borne. Using these strict criteria, we analyzed the literature and identified one strongly supported occurrence of CRISPR-Cas compromise. The MDR *E. faecalis* strain KUB3006, for which a completely closed genome is available (44), possesses 4 plasmids, including one that encodes linezolid resistance. Yet, it also encodes CRISPR3-Cas (44). The *cas9* gene is frameshifted, and the frameshift occurs within the codon for one of the two Cas9 active sites that we previously experimentally confirmed in *E. faecalis* (17). We conclude that CRISPR3-Cas function is likely to be compromised in this strain, which requires experimental confirmation.

## Discussion

In this study we investigated the fates of *E. faecalis* transconjugants that acquired a CRISPR-targeted plasmid, using serial passage with and without antibiotic selection for the plasmid.

We observed that 5 of 6 wild-type T11RF(pAM714) transconjugants lost the plasmid over the course of antibiotic-free serial passage, while T11RFΔ*cas9*(pAM714) transconjugants maintained the plasmid. This is indicative of active CRISPR-Cas defense in the T11RF(pAM714) transconjugant populations. An important caveat is that erythromycin is a bacteriostatic antibiotic, and we did not restreak transconjugant colonies before starting the serial passage experiment, therefore we cannot be certain that plasmid-free T11RF surviving on the transconjugant selective agar were not carried over with the initial transconjugant colonies that were picked. At least two non-mutually exclusive explanations for our observations are possible. The first is that T11RF cells lacking pAM714 (either carried over or sporadically arising) outcompete T11RF(pAM714) cells. This competitive effect is due to the growth defect of cells that simultaneously possess CRISPR-Cas and one of its plasmid targets, as we have previously reported (15, 16). The second is that CRISPR-Cas ‘scrubs’ pAM714 from T11RF over the course of antibiotic-free serial passage. We point to Figure S1, where the WT4 colony stands out as containing essentially all erythromycin-resistant cells, while the other WT transconjugant colonies do not. As determined in this study, WT4 is a *cas9* loss-of-function mutant. These results seem to favor the second explanation, since carryover of erythromycin-sensitive cells may have equally affected the randomly selected WT4 colony compared to other randomly selected WT colonies. Regardless, this could be further investigated by repeating the experiment and streaking the initial transconjugant colonies prior to serial passage.

On the other hand, when the same T11RF(pAM714) transconjugants were passaged with antibiotic selection, the plasmid was maintained, as expected, but variants emerged that lacked the spacer targeting the plasmid. In some cases, multiple sequential spacers were deleted, and we demonstrated that this resulted in loss of defense against multiple plasmids (specifically, pAM714 and derivatives of pCF10; see WT5, Fig. 3). This supports our CRISPR amplicon sequence analysis by demonstrating that spacer losses identified by sequencing corresponded with loss of functional CRISPR-Cas defense. Variations in *cas9* were also observed in some antibiotic-passaged populations, but not experimentally investigated, so their impacts on CRISPR-Cas function remain undefined. We also identified one very strongly supported example of CRISPR-Cas compromise in an independent *E. faecalis* isolate (KUB3006) with a closed genome. Other studies have identified spacer deletion events occurring in transformants or transconjugants of CRISPR-targeted plasmids (45–47) providing more evidence that loss-of-function mutations in CRISPR-Cas contribute to antibiotic resistance plasmid dissemination in bacterial populations.

Our work occurs in the broader context of work on “self-targeting” spacers in CRISPR-Cas systems, which have been studied for some time across different types of CRISPR-Cas in different species (48). Self-targeting spacers typically refer to those with sequence complementarity to targets within the host genome. This phenomenon was mostly observed during phage infection and plasmid invasion (49), but self-targeting spacers against non-mobile elements have also been reported (50). Different hypotheses have been posited for their maintenance in microbial genomes, including alteration of target sites via DNA repair (51–53), large deletion of the target sites (54), alteration of the PAM sequence (55), loss of spacers (16, 56) or mutations in *cas* genes (54). The “self-targeting” outcome is dependent on the fitness cost and environmental selective pressure (57). The conflict between active CRISPR-Cas and “self-targeting” spacers plays a role in shaping bacterial evolution, including altering the population-level genetic diversity (58–60), remodeling of pathogenicity islands (61) and modulation of metabolic pathways (62). Our work sought to investigate how this conflict can be resolved in *E. faecalis*, as entry and maintenance of pAM714 into wild-type T11RF creates a “self-targeting” situation that theoretically must be resolved to prevent persistent stress from DNA damage at the targeted pAM714 *repB* site. In essence, this is the basis of CRISPR-Cas gene editing, and in a previous study we took advantage of this property to implement CRISPR-Cas9 gene editing in *E. faecalis* (15). Here, a recombination template to repair DNA damage (in this case, damage to pAM714 *repB*) was not provided to the cells. We observed (in general) that the “self-targeting” CRISPR spacer was retained and the target lost in populations passaged without antibiotic selection for the target. On the other hand, CRISPR spacer and/or potentially overall CRISPR-Cas function was lost, and the target retained, in populations passaged with antibiotic selection for the target. In other studies, these “self-targeting” spacers have been engineered to promote the loss of genomic islands and other mobile elements in bacteria (54, 61, 63). Our results demonstrate that selection for the targeted element impedes this process, which is relevant to the design and implementation of CRISPR-based antimicrobials that “cure” *E. faecalis* of antibiotic resistance genes (19, 64). Our results suggest that these systems will need to be introduced and utilized either pre- or post-antibiotic therapy, not during.

Overall, we posit that the interplay of CRISPR-Cas, plasmids, and antibiotic selection should be further investigated to understand the role of CRISPR-Cas in the antibiotic resistance crisis. Particularly important will be experimental designs that better replicate the *in vivo* setting where antibiotic resistance plasmids disseminate, for e.g., the gastrointestinal tract, where multispecies biofilms are present, and cell densities, mutation rates, CRISPR-Cas activities, and antibiotic concentrations are likely to vary. More specifically, it remains unclear how frequently CRISPR-Cas spacer deletion or loss-of-function mutants arise *in vivo* under antibiotic selection for “self-targeted” plasmids. In a prior study, we assessed the *in vivo* functioning of T11RF CRISPR-Cas against pAM714 in a mouse intestinal colonization model where OG1SSp(pAM714) donors and T11RF or T11RFΔ*cas9* recipients were introduced (18). CRISPR-Cas defense against pAM714 appeared robust – no T11RF(pAM714) transconjugants stably colonized the mouse intestine above our limit of detection (<10^2^ CFU/g feces). Conversely, T11RFΔ*cas9*(pAM714) transconjugants were stably present in most mice (up to ∼10^7^ CFU/g feces). The key question we did not answer in that study is whether T11RF(pAM714) transconjugants with spacer 6 deletions or other loss-of-function mutations in CRISPR-Cas would have emerged if we had treated the mice with erythromycin. This remains an outstanding question to be addressed.

## Materials and Methods

### Strains, reagents, and routine molecular biology procedures

Bacterial strains and plasmids used in this study are listed in Table 1. *E. faecalis* strains were grown in Brain Heart Infusion (BHI) broth or on agar plates at 37°C unless otherwise noted. Antibiotics were used for *E. faecalis* at the following concentrations: erythromycin, 50 μg/mL; chloramphenicol, 15 μg/mL; streptomycin, 500 μg/mL; spectinomycin, 500 μg/mL; rifampicin, 50 μg/mL; fusidic acid, 25 μg/mL; tetracycline, 10 μg/mL. *Escherichia coli* strains used for plasmid propagation and were grown in lysogeny broth (LB) broth or on agar plates at 37°C. Chloramphenicol was used at 15 μg/mL for *E. coli*. PCR was performed using *Taq* (New England Biolabs) or Phusion (Fisher Scientific) polymerases. Primer sequences used are in Table S1. Routine Sanger sequencing was carried out at the Massachusetts General Hospital DNA core facility (Boston, MA). *E. faecalis* electrocompetent cells were made using the lysozyme method as previously described (65).

### Generation of pCF10 derivatives

To insert the T11 CRISPR3 spacer 1 (S_1_), S_6_, and S_7_ sequences and CRISPR3 PAM (TTGTA) into pCF10, 47 bp and 39 bp single stranded DNA oligos were annealed to each other to generate dsDNA with restriction enzyme overhangs for BamHI and PstI. The annealed oligos were ligated into the pLT06 derivative pWH107 that includes sequence from pCF10 *uvrB*, to insert these sequences into the *uvrB* gene of pCF10 by homologous recombination. A knock-in protocol was performed as previously described (17).

### Conjugation experiments

*E. faecalis* donor and recipient strains were grown in BHI overnight to stationary phase. A 1:10 dilution was made for both donor and recipient cultures in fresh BHI broth and incubated for 1.5 hr to reach mid-exponential phase. A mixture of 100 μL donor cells and 900 μL recipient cells was pelleted and plated on non-selective BHI agar to allow conjugation. After 18 h incubation, the conjugation mixture was scraped from the plate using 2 mL 1X PBS supplemented with 2 mM EDTA. Serial dilutions were prepared from the conjugation mixture and plated on BHI agars selective for transconjugants or donors. After 24-48 h incubation, colony forming units per milliliter (CFU/mL) was determined using plates with 30 - 300 colonies. The conjugation frequency was calculated as the CFU/mL of transconjugants divided by the CFU/mL of donors.

### Serial passage

Transconjugant colonies were suspended in 50 μL BHI broth. The 50 μL suspension was used as follows: 3 μL was used for PCR to confirm the integrity of the CRISPR array, 10 μL was inoculated into plain BHI broth, another 10 μL was inoculated into selective BHI broth for plasmid selection, and another 10 μL was used for serial dilution and plating on selective medium to enumerate the initial number of plasmid-containing cells in the transconjugant colonies. Broth cultures were incubated for 24 h, followed by 1:1000 dilution into either fresh plain BHI or fresh selective BHI. At each 24 h interval, 3 μL of each culture from the previous incubation was used for PCR to check CRISPR array integrity, and 10 μL was used for serial dilution and plating on agars to determine CFU/mL for total viable cells and plasmid-containing cells. The cultures were passaged in this manner for 14 days; cryopreserved culture stocks were made daily in glycerol. To use the Day 14 transconjugant populations in conjugation reactions, the glycerol stocks were completely thawed on ice, and 20 μL was inoculated into plain BHI broth. The cultures were incubated for 6-8 h to allow them to reach mid-exponential phase (OD_600nm_ ≈ 0.5–0.7), and 900 μL was used as recipient in conjugation reactions as described above.

### Deep sequencing of CRISPR3 amplicons and genomic DNA

For CRISPR3 amplicon sequencing, 3 μL from a broth culture was used as template in PCR using Phusion Polymerase with CR3_seq_F/R primers (Table S1). The PCR products were purified using the Thermo Scientific PCR purification kit (Thermo Scientific). Genomic DNA was isolated using the phenol-chloroform method (66). The purified PCR amplicons and genomic DNA samples were sequenced using 2 x 150 bp paired end sequencing chemistry by Molecular Research LP (MR DNA; Texas).

### Whole genome sequencing analysis

T11 supercontig and pAD1 plasmid contig references were downloaded from NCBI (accession numbers: T11: NZ_GG688637.1-NZ_GG688649; pAD1: AB007844, AF394225, AH011360, L01794, L19532, L37110, M84374, M87836, U00681, X17214, X62657, X62658, M11180 [Tn*917*]). Reads were aligned to these references using default parameters in CLC Genomics Workbench (Qiagen) where ≥50% of each mapped read has ≥80% sequence identity to the reference. Variations occurring with ≥35% frequency at positions with ≥10X coverage between our samples and the reference contigs were detected using the Basic Variant Detector. At the same time, local realignment was performed, followed by Fixed Ploidy variant detection using default parameters and variants probability ≥90% in CLC Genomics Workbench. The basic variants and fixed ploidy variants were combined for each sequencing sample and subjected to manual inspection. The variants that were detected in the T11 genome from all samples were inferred to be variants in our parent T11 stock and were manually removed. The variants that were detected in pAD1 genome from all transconjugant samples were inferred to be variants in our pAM714 stock, hence were also manually removed. Next, variants within the CRISPR3 array were removed as we analyzed CRISPR3 alleles using a different approach (amplicon deep sequencing; see below). All variants detected from all populations were manually checked for coverage depth to eliminate the detection bias. The variants detected in all samples are shown in Table S2. To detect the insertion site of Tn*917*, the mapped reads on reference M11180 were inspected. Reads immediately adjacent to the 5’ and 3’ ends of Tn*917* shared consensus sequences which were later used for blast against T11 and pAD1 (Figure S5). No evidence for Tn*917* insertions other than in the expected position in pAM714 was detected.

### Analysis of CRISPR3 amplicon sequencing

To analyze the amplicon sequencing, we first created a pool of references with 96-bp in length to represent both wild type and mutant CRISPR3 alleles. In the wild type reference pool, each 96-bp reference contains two adjacent spacers connected by one direct repeat (5’-S_x_-R-S_x+1_-3’ where x is from 1 to 20). To include the leader end and terminal end, wild type reference pool also contains 30 bp upstream of the first repeat (S_0_) and 30 bp downstream of the terminal repeat (S_T_), generating two additional wild type references: 5’-S_0_-R-S_1_-3’ and 5’-S_21_-TR-S_T_-3’. In total, 22 references were created to represent wild type CRISPR3 allele.

To detect CRISPR3 spacer deletions and rearrangements, we manually created an additional 462 references with 96-bp in length to represent mutant CRISPR3 alleles. Each mutant reference contains two spacers connected by a direct repeat, where the reference was not already represented in the wild type reference pool. Schematically, the mutant references can be expressed by 5’-spacer[x]-repeat-spacer[y]-3’ (5’-S_x_RS_y_-3’) where y ≠ x. The selection of spacer[x] is from S_0_ to S_21_ while the selection of spacer[y] is from S_1_ to S_T_. Additionally, we assume that if spacer[y] is ST then the repeat would be terminal repeat TR, generating 5’-S_x_TRS_T_-3’ (x is from 0 to 21). In total, 462 references were created with 96-bp in length.

Sequencing reads were first clipped into 75 bp fragments to enhance mapping efficiency, allowing for retrieval of maximal sequence information. The clipped reads were used to map to the pool of 96-bp references using stringent mapping parameters in CLC Genomics Workbench (Qiagen). The stringent mapping parameters require 100% of each mapped read to be ≥95% identical to one unique reference. Thus, the sequencing reads from different CRISPR alleles will be distinguished. The number of mapped reads per each reference was calculated and normalized by the total number of mapped reads to all references, generating normalized reads per reference. The total number of mapped reads to pools of wild type references and mutant references were summarized in Table S3.

To visualize the normalized reads per reference, a heatmap was generated (Figure S4). On the heatmap, each row name represents 5’-S_x_ (x is from 0 to 20) and each column name represents R-S_y_-3’ (y is from 1 to 21) or TR-S_T_-3’. The diagonal cells are empty due to the assumption of no repetitive spacers (or 5’-Sx-R-Sx-3’). Intuitively, the cells adjacent to the diagonal cells on the upper side represents wild type pool of references, 5’-Sx-R-Sy-3’ where y=x+1 (Figure S3). The cell color represents normalized reads per reference (from white to red to black is lowest to intermediate to the highest abundance).

To evaluate the statistical significance of normalized mapped reads, we calculated z-score using the mean and standard deviation calculated from T11RF control. Combining the normalized reads per reference from Day 1 and Day 14 T11RF control, we obtained an average normalized reads per reference of 0.002 or 2%) with standard deviation of 0.0096 or 9.6%. These numbers were used to calculated z-score for normalized reads per reference for all samples. To assess the significance, the z-score is further transformed into p-value. A p-value of 0.05 was used as a significant cutoff.

To visualize the p-value and derive mutant alleles, a heatmap was plotted using the same method as above. The cells with p-value < 0.05 were color coded. To derive the mutant allele, we assume that S_0_ and S_T_ are intact. The sequence of spacers was derived based on the idea of forward algorithm and Viterbi algorithm. Each p-value was considered as a conditional probability: p-value (5’-Sx-R-Sy-3’) ∼ P(spacer y |_spacer x_).

### Accession number

The sequencing data for amplicon and whole genome sequencing analysis of transconjugant populations has been deposited in the NCBI Sequence Read Archive under PRJNA418345.

## Acknowledgement

This work was supported by Public Health Service grant R01AI116610 to K.L.P and the Cecil H. and Ida Green Chair to M.Q.Z. We thank Dr. Chen Jia for consultation on data analysis methods.

---

**Figure S1.**
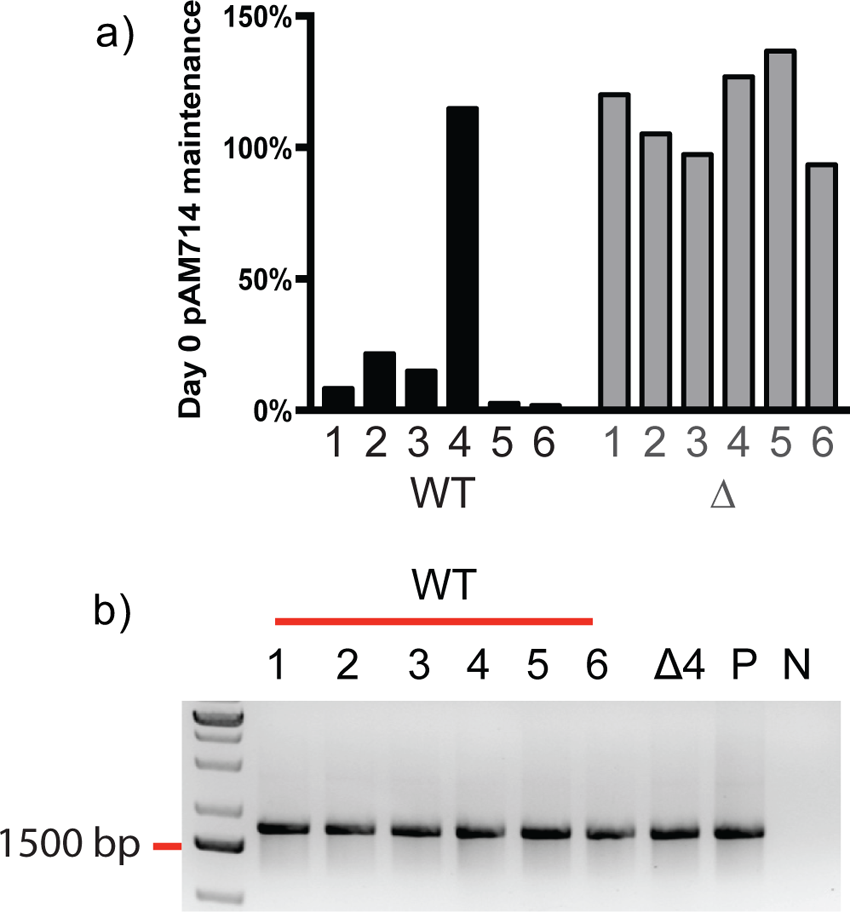
Initial phenotypes of transconjugants chosen for serial passage experiments. a) Frequency of erythromycin resistance in transconjugant colonies used to initiate serial passage experiments. Erythromycin resistance is used to track pAM714 maintenance in the population. b) CRISPR3 amplicon PCR results for transconjugant colonies used to initiate serial passage experiments. Shown are CRISPR3 amplicon sizes for six T11RF pAM714 transconjugants (WT 1-6), a representative T11RFΔ*cas9* pAM714 transconjugant (Δ4), T11RF genomic DNA as a positive control (P), and a reagent control (N).

**Figure S2.**
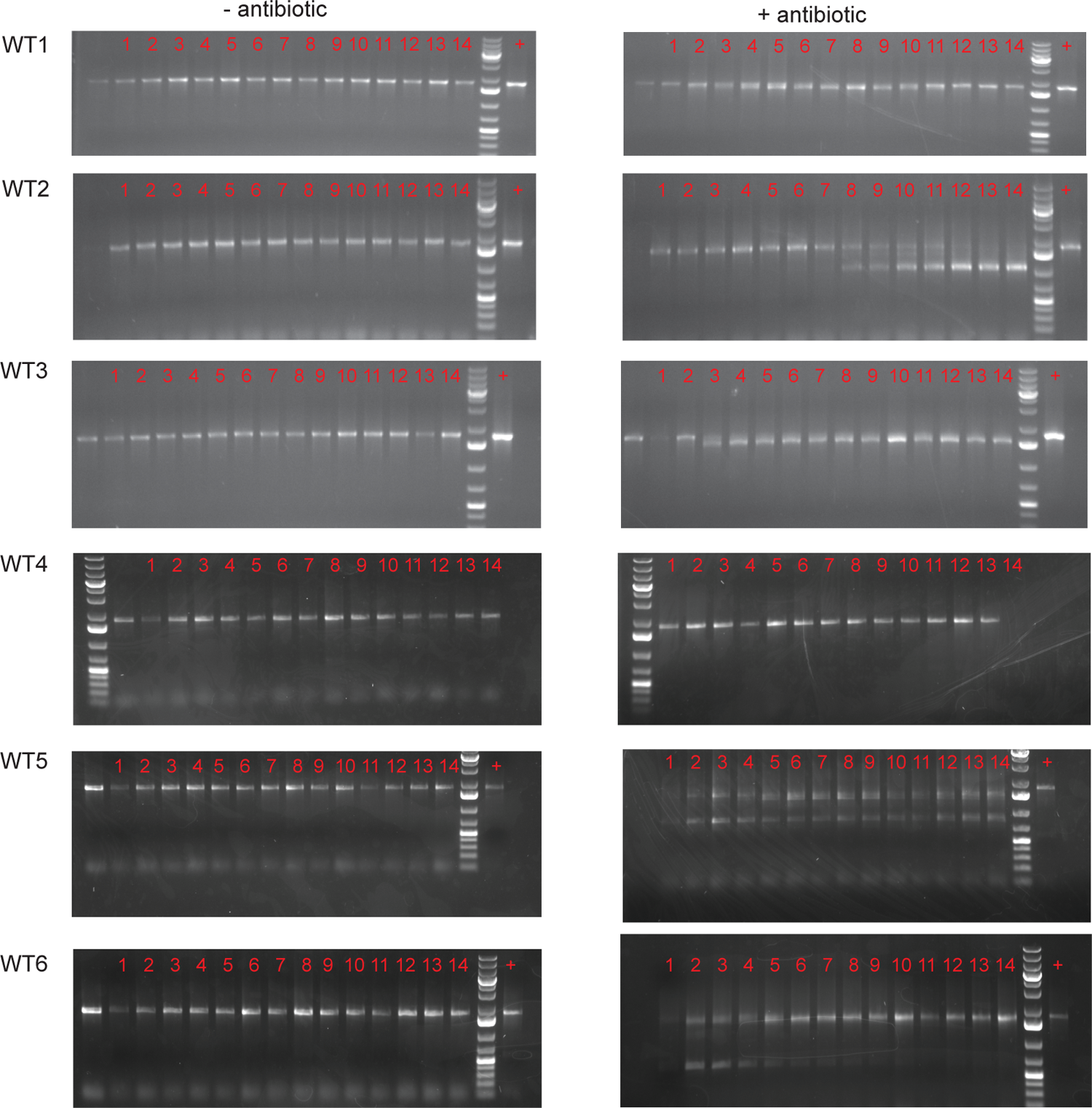
Agarose gel electrophoresis analysis of CRISPR3 amplicons in six T11RF(pAM714) transconjugant populations. Six T11RF(pAM714) transconjugants were serially passaged for 14 days in BHI (left panel) or BHI with erythromycin (right panel). The size of the CRISPR3 array was monitored using PCR and gel electrophoresis on each passage day.

**Figure S3.**
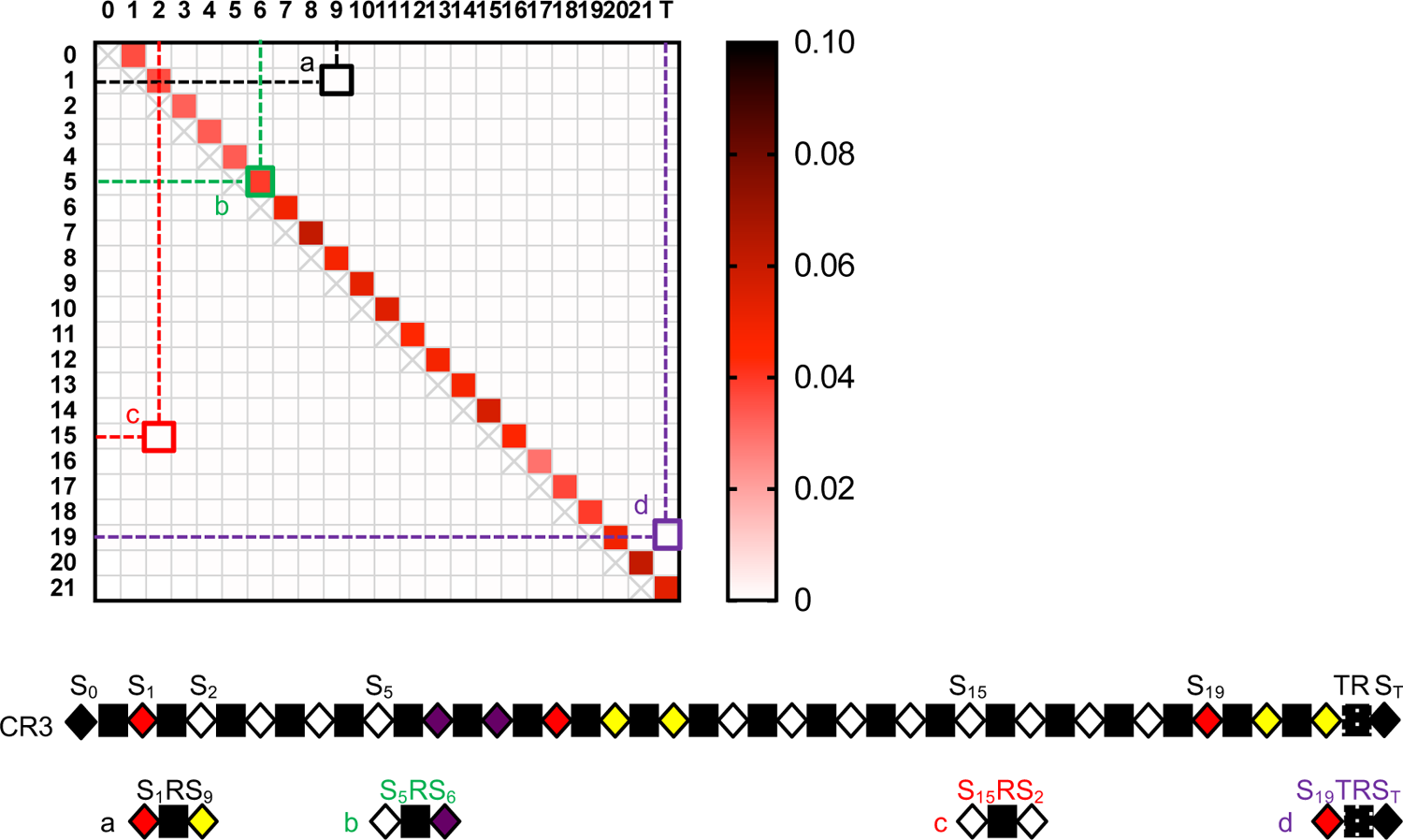
Demonstration of mutant alleles and visualization of heatmap matrix. A cartoon of the T11 CRISPR3 array is shown below the heatmap. The array contains 21 unique spacer sequences, of which spacer 6 targets pAM714. Four scenarios (a-d) corresponding to four cells indicating different wild-type (b) or mutant (a,c,d) allele types in the matrix are shown.

**Figure S4.**
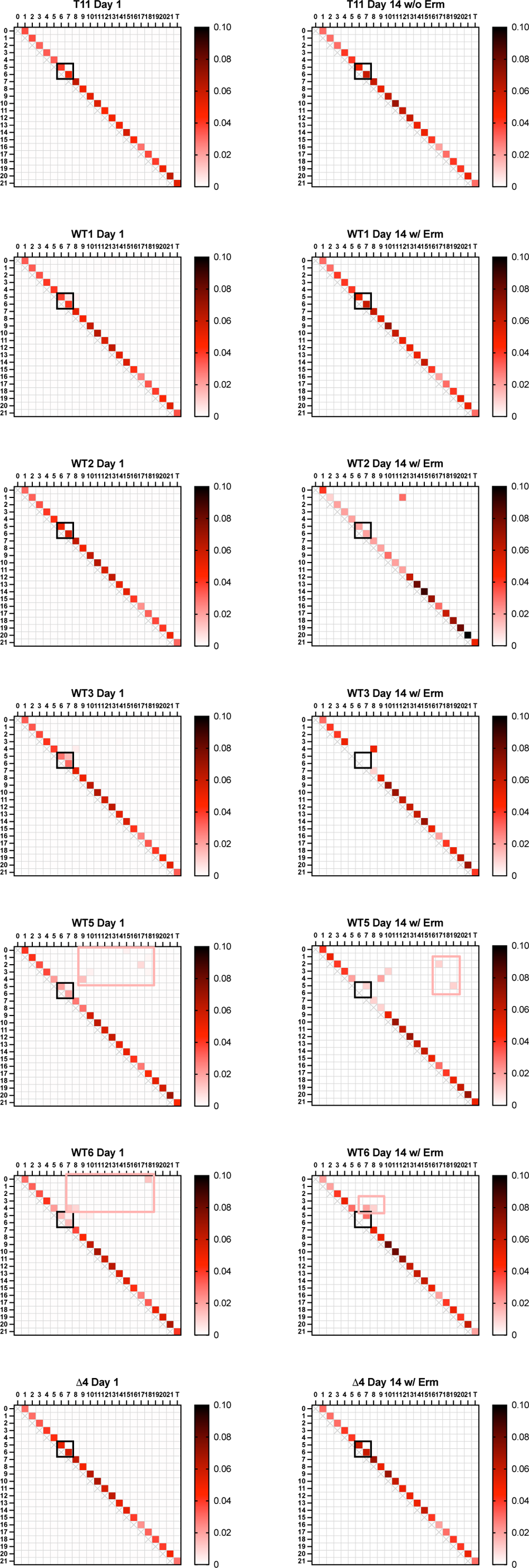
Less abundant mutant alleles can be observed in a heatmap of normalized read counts. Heat maps show the abundance of each mutant allele per sample. Day 1 and Day 14 populations were analyzed and T11RF was used as a control. Each cell within the heat map represents one of the 484 artificial references. The color scale represents low (white) to intermediate (ref) to high (black) abundance alleles. Black boxes indicate the expected wild type allele at the S5-S6 region. Pink boxes surround rare alleles that did not meet the significance cutoff but can be visible here.

**Figure S5.**
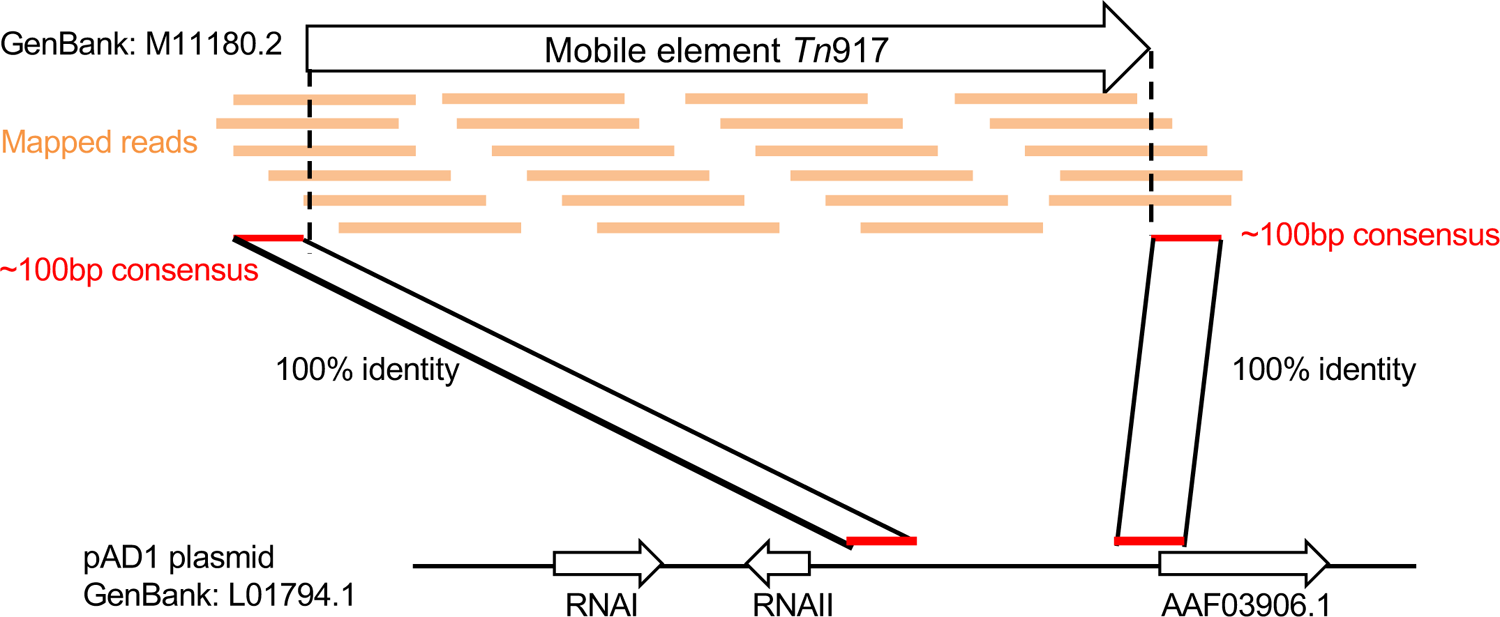
Mapping of Tn*917* location. The consensus sequences were extracted from 5’ and 3’ ends of transposon Tn*917* and mapped to pAD1 sequences.

**Table S1.**
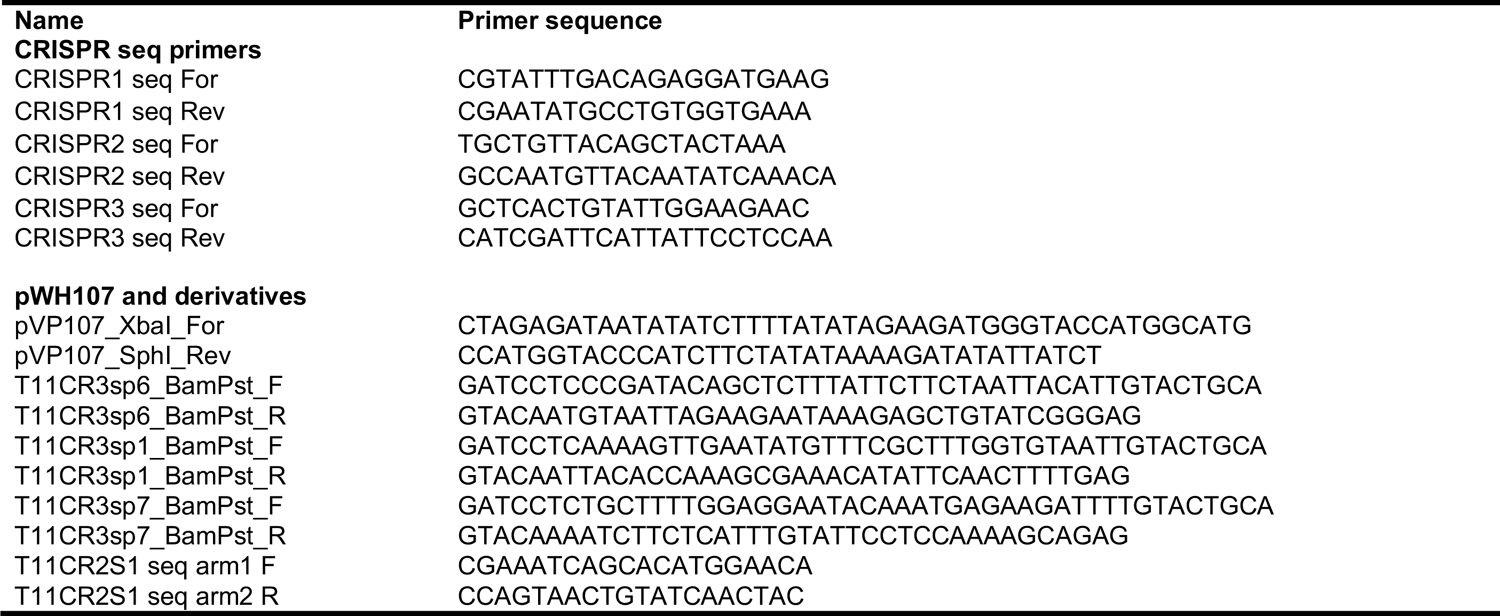
Primers used in this study.

**Table S2.**
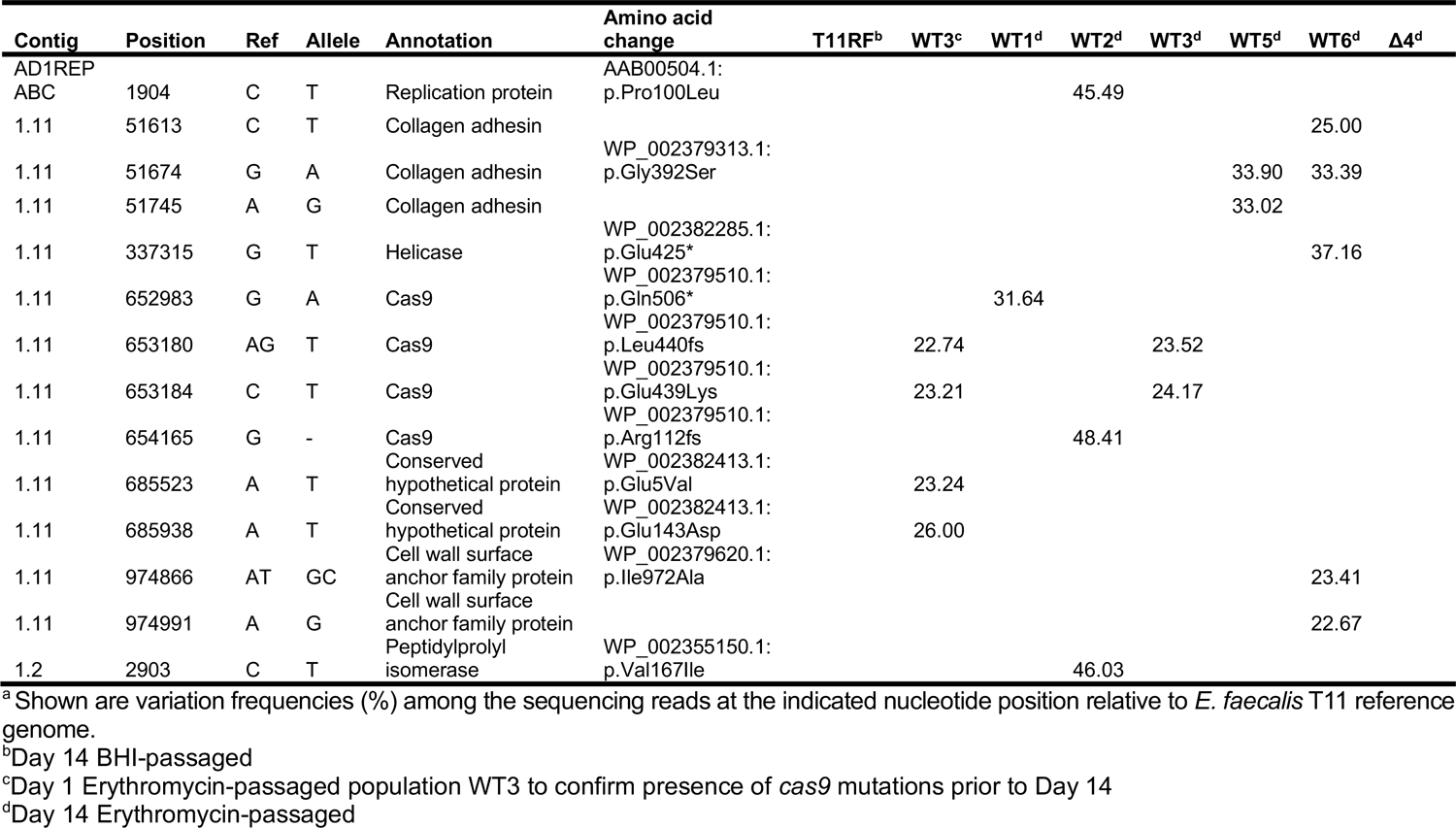
SNP detection in all genomic DNA sequencing samples.^a^

**Table S3.** Mapping statistics of the amplicon sequencing.

| **Sample name** | **# of Total reads** | **# mapped to WT pool of references** | **# mapped to mutant pool of references** |
| --- | --- | --- | --- |
| **Day 1** |  |  |  |
| T11RF | 19,082,652 | 7,273,291 | 125,204 |
| Δ4 | 20,808,404 | 8,995,910 | 77,551 |
| WT1 | 16,736,162 | 6,808,760 | 168,051 |
| WT2 | 16,859,152 | 7,222,339 | 59,609 |
| WT3 | 17,357,970 | 7,136,664 | 250,783 |
| WT5 | 17,744,230 | 6,784,881 | 430,500 |
| WT6 | 16,179,400 | 6,337,706 | 406,347 |
| **Day 14** |  |  |  |
| T11RF | 17,973,946 | 7,760,594 | 68,900 |
| Δ4 | 14,403,868 | 5,369,063 | 55,037 |
| WT1 | 14,158,158 | 5,219,558 | 55,730 |
| WT2 | 14,538,836 | 4,489,642 | 254,615 |
| WT3 | 17,174,946 | 5,653,212 | 385,906 |
| WT5 | 15,200,646 | 5,384,356 | 460,061 |
| WT6 | 12,325,042 | 4,201,804 | 303,973 |

## References

1. Lebreton F, Willems RJL, Gilmore MS. 2014. Enterococcus Diversity, Origins in Nature, and Gut Colonization. In Gilmore MS, Clewell DB, Ike Y, Shankar N (ed), Enterococci: From Commensals to Leading Causes of Drug Resistant Infection, Boston.

2. Kristich CJ, Djoric D, Little JL. 2014. Genetic basis for vancomycin-enhanced cephalosporin susceptibility in vancomycin-resistant enterococci revealed using counterselection with dominant-negative thymidylate synthase. Antimicrob Agents Chemother 58:1556–64. 3957902

3. Uttley AH, George RC, Naidoo J, Woodford N, Johnson AP, Collins CH, Morrison D, Gilfillan AJ, Fitch LE, Heptonstall J. 1989. High-level vancomycin-resistant enterococci causing hospital infections. Epidemiol Infect 103:173–81. 2249484

4. Huycke MM, Sahm DF, Gilmore MS. 1998. Multiple-drug resistant enterococci: the nature of the problem and an agenda for the future. Emerging infectious diseases 4:239.

5. Clewell DB, Weaver KE, Dunny GM, Coque TM, Francia MV, Hayes F. 2014. Extrachromosomal and Mobile Elements in Enterococci: Transmission, Maintenance, and Epidemiology. In Gilmore MS, Clewell DB, Ike Y, Shankar N (ed), Enterococci: From Commensals to Leading Causes of Drug Resistant Infection, Boston.

6. Palmer KL, Kos VN, Gilmore MS. 2010. Horizontal gene transfer and the genomics of enterococcal antibiotic resistance. Curr Opin Microbiol 13:632–9. 2955785

7. Hegstad K, Mikalsen T, Coque T, Werner G, Sundsfjord A. 2010. Mobile genetic elements and their contribution to the emergence of antimicrobial resistant *Enterococcus faecalis* and *Enterococcus faecium*. Clinical microbiology and infection 16:541–554.

8. Clewell DB. 2007. Properties of *Enterococcus faecalis* plasmid pAD1, a member of a widely disseminated family of pheromone-responding, conjugative, virulence elements encoding cytolysin. Plasmid 58:205–27.

9. Dunny GM. 2007. The peptide pheromone-inducible conjugation system of *Enterococcus faecalis* plasmid pCF10: cell-cell signalling, gene transfer, complexity and evolution. Philos Trans R Soc Lond B Biol Sci 362:1185–93. 2435581

10. Gilmore MS, Segarra RA, Booth MC, Bogie CP, Hall LR, Clewell DB. 1994. Genetic structure of the Enterococcus faecalis plasmid pAD1-encoded cytolytic toxin system and its relationship to lantibiotic determinants. J Bacteriol 176:7335–44. PMC197123

11. Jasni AS, Mullany P, Hussain H, Roberts AP. 2010. Demonstration of conjugative transposon (Tn*5397*)-mediated horizontal gene transfer between *Clostridium difficile* and *Enterococcus faecalis*. Antimicrob Agents Chemother 54:4924–6. 2976158

12. Showsh SA, De Boever EH, Clewell DB. 2001. Vancomycin Resistance Plasmid in *Enterococcus faecalis* That Encodes Sensitivity to a Sex Pheromone Also Produced by *Staphylococcus aureus*. Antimicrobial Agents and Chemotherapy 45:2177–2178.

13. Tsvetkova K, Marvaud JC, Lambert T. 2010. Analysis of the mobilization functions of the vancomycin resistance transposon Tn1549, a member of a new family of conjugative elements. J Bacteriol 192:702–13. 2812457

14. Zhu W, Murray PR, Huskins WC, Jernigan JA, McDonald LC, Clark NC, Anderson KF, McDougal LK, Hageman JC, Olsen-Rasmussen M, Frace M, Alangaden GJ, Chenoweth C, Zervos MJ, Robinson-Dunn B, Schreckenberger PC, Reller LB, Rudrik JT, Patel JB. 2010. Dissemination of an *Enterococcus* Inc18-Like *vanA* plasmid associated with vancomycin-resistant *Staphylococcus aureus*. Antimicrob Agents Chemother 54:4314–20. 2944587

15. Hullahalli K, Rodrigues M, Nguyen UT, Palmer K. 2018. An Attenuated CRISPR-Cas System in *Enterococcus faecalis* Permits DNA Acquisition. MBio 9:e00414–18. PMC5930301

16. Hullahalli K, Rodrigues M, Palmer KL. 2017. Exploiting CRISPR-Cas to manipulate *Enterococcus faecalis* populations. Elife 6:e26664. PMC5491264

17. Price VJ, Huo W, Sharifi A, Palmer KL. 2016. CRISPR-Cas and Restriction-Modification Act Additively against Conjugative Antibiotic Resistance Plasmid Transfer in Enterococcus faecalis. mSphere 1:e00064-16.

18. Price VJ, McBride SW, Hullahalli K, Chatterjee A, Duerkop BA, Palmer KL. 2019. Enterococcus faecalis CRISPR-Cas Is a Robust Barrier to Conjugative Antibiotic Resistance Dissemination in the Murine Intestine. mSphere 4:e00464-19. PMC6656873

19. Rodrigues M, McBride SW, Hullahalli K, Palmer KL, Duerkop BA. 2019. Conjugative Delivery of CRISPR-Cas9 for the Selective Depletion of Antibiotic-Resistant Enterococci. Antimicrob Agents Chemother 63:e01454–19. PMC6811441

20. Barrangou R, Fremaux C, Deveau H, Richards M, Boyaval P, Moineau S, Romero DA, Horvath P. 2007. CRISPR provides acquired resistance against viruses in prokaryotes. Science 315:1709–12.

21. Marraffini LA, Sontheimer EJ. 2008. CRISPR interference limits horizontal gene transfer in staphylococci by targeting DNA. Science 322:1843–5. 2695655

22. Bourgogne A, Garsin DA, Qin X, Singh KV, Sillanpaa J, Yerrapragada S, Ding Y, Dugan-Rocha S, Buhay C, Shen H, Chen G, Williams G, Muzny D, Maadani A, Fox KA, Gioia J, Chen L, Shang Y, Arias CA, Nallapareddy SR, Zhao M, Prakash VP, Chowdhury S, Jiang H, Gibbs RA, Murray BE, Highlander SK, Weinstock GM. 2008. Large scale variation in *Enterococcus faecalis* illustrated by the genome analysis of strain OG1RF. Genome Biol 9:R110. 2530867

23. Hullahalli K, Rodrigues M, Schmidt BD, Li X, Bhardwaj P, Palmer KL. 2015. Comparative Analysis of the Orphan CRISPR2 Locus in 242 *Enterococcus faecalis* Strains. PLoS One 10:e0138890. 4580645

24. Palmer KL, Gilmore MS. 2010. Multidrug-resistant enterococci lack CRISPR-*cas*. MBio 1. 2975353

25. Garneau JE, Dupuis ME, Villion M, Romero DA, Barrangou R, Boyaval P, Fremaux C, Horvath P, Magadan AH, Moineau S. 2010. The CRISPR/Cas bacterial immune system cleaves bacteriophage and plasmid DNA. Nature 468:67–71.

26. Horvath P, Romero DA, Coute-Monvoisin AC, Richards M, Deveau H, Moineau S, Boyaval P, Fremaux C, Barrangou R. 2008. Diversity, activity, and evolution of CRISPR loci in *Streptococcus thermophilus*. J Bacteriol 190:1401–12. 2238196

27. Deveau H, Barrangou R, Garneau JE, Labonte J, Fremaux C, Boyaval P, Romero DA, Horvath P, Moineau S. 2008. Phage response to CRISPR-encoded resistance in *Streptococcus thermophilus*. J Bacteriol 190:1390–400. 2238228

28. Mojica FJ, Diez-Villasenor C, Garcia-Martinez J, Almendros C. 2009. Short motif sequences determine the targets of the prokaryotic CRISPR defence system. Microbiology 155:733–40.

29. Yosef I, Goren MG, Qimron U. 2012. Proteins and DNA elements essential for the CRISPR adaptation process in *Escherichia coli*. Nucleic Acids Res 40:5569–76. 3384332

30. Nunez JK, Kranzusch PJ, Noeske J, Wright AV, Davies CW, Doudna JA. 2014. Cas1-Cas2 complex formation mediates spacer acquisition during CRISPR-Cas adaptive immunity. Nat Struct Mol Biol doi:10.1038/nsmb.2820.

31. Heler R, Samai P, Modell JW, Weiner C, Goldberg GW, Bikard D, Marraffini LA. 2015. Cas9 specifies functional viral targets during CRISPR–Cas adaptation. Nature 519:199.

32. Deltcheva E, Chylinski K, Sharma CM, Gonzales K, Chao Y, Pirzada ZA, Eckert MR, Vogel J, Charpentier E. 2011. CRISPR RNA maturation by trans-encoded small RNA and host factor RNase III. Nature 471:602–7. 3070239

33. Jinek M, Jiang F, Taylor DW, Sternberg SH, Kaya E, Ma E, Anders C, Hauer M, Zhou K, Lin S, Kaplan M, Iavarone AT, Charpentier E, Nogales E, Doudna JA. 2014. Structures of Cas9 endonucleases reveal RNA-mediated conformational activation. Science 343:1247997.

34. Sternberg SH, Redding S, Jinek M, Greene EC, Doudna JA. 2014. DNA interrogation by the CRISPR RNA-guided endonuclease Cas9. Nature 507:62–7.

35. Jinek M, Chylinski K, Fonfara I, Hauer M, Doudna JA, Charpentier E. 2012. A programmable dual-RNA-guided DNA endonuclease in adaptive bacterial immunity. Science 337:816–21.

36. Gasiunas G, Barrangou R, Horvath P, Siksnys V. 2012. Cas9-crRNA ribonucleoprotein complex mediates specific DNA cleavage for adaptive immunity in bacteria. Proc Natl Acad Sci U S A 109:E2579–86. 3465414

37. Huo W, Price VJ, Sharifi A, Zhang MQ, Palmer KL. 2018. Enterococcus faecalis strains with compromised CRISPR-Cas defense emerge under antibiotic selection for a CRISPR-targeted plasmid. bioRxiv doi:10.1101/220467:220467.

38. McBride SM, Fischetti VA, Leblanc DJ, Moellering RC, Jr., Gilmore MS. 2007. Genetic diversity among *Enterococcus faecalis*. PLoS One 2:e582. 1899230

39. Palmer KL, Godfrey P, Griggs A, Kos VN, Zucker J, Desjardins C, Cerqueira G, Gevers D, Walker S, Wortman J, Feldgarden M, Haas B, Birren B, Gilmore MS. 2012. Comparative genomics of enterococci: variation in *Enterococcus faecalis*, clade structure in *E. faecium*, and defining characteristics of *E. gallinarum* and *E. casseliflavus*. MBio 3:e00318–11. 3374389

40. Weaver KE, Tritle DJ. 1994. Identification and characterization of an *Enterococcus faecalis* plasmid pAD1-encoded stability determinant which produces two small RNA molecules necessary for its function. Plasmid 32:168–81.

41. Ike Y, Clewell DB, Segarra RA, Gilmore MS. 1990. Genetic analysis of the pAD1 hemolysin/bacteriocin determinant in *Enterococcus faecalis*: Tn*917* insertional mutagenesis and cloning. J Bacteriol 172:155–63. 208413

42. Clewell DB, Tomich PK, Gawron-Burke MC, Franke AE, Yagi Y, An FY. 1982. Mapping of *Streptococcus faecalis* plasmids pAD1 and pAD2 and studies relating to transposition of Tn*917*. J Bacteriol 152:1220–30. 221629

43. Dunny G, Yuhasz M, Ehrenfeld E. 1982. Genetic and physiological analysis of conjugation in Streptococcus faecalis. Journal of bacteriology 151:855–859.

44. Kuroda M, Sekizuka T, Matsui H, Suzuki K, Seki H, Saito M, Hanaki H. 2018. Complete Genome Sequence and Characterization of Linezolid-Resistant *Enterococcus faecalis* Clinical Isolate KUB3006 Carrying a *cfr*(B)-Transposon on Its Chromosome and *optrA*-Plasmid. Front Microbiol 9:2576. PMC6209644

45. Jiang W, Maniv I, Arain F, Wang Y, Levin BR, Marraffini LA. 2013. Dealing with the evolutionary downside of CRISPR immunity: bacteria and beneficial plasmids. PLoS Genet 9:e1003844. 3784566

46. Rao C, Chin D, Ensminger AW. 2017. Priming in a permissive type I-C CRISPR-Cas system reveals distinct dynamics of spacer acquisition and loss. RNA 23:1525–1538. PMC5602111

47. Lopez-Sanchez MJ, Sauvage E, Da Cunha V, Clermont D, Ratsima Hariniaina E, Gonzalez-Zorn B, Poyart C, Rosinski-Chupin I, Glaser P. 2012. The highly dynamic CRISPR1 system of *Streptococcus agalactiae* controls the diversity of its mobilome. Mol Microbiol 85:1057–71.

48. Wimmer F, Beisel CL. 2019. CRISPR-Cas Systems and the Paradox of Self-Targeting Spacers. Front Microbiol 10:3078. PMC6990116

49. Staals RH, Jackson SA, Biswas A, Brouns SJ, Brown CM, Fineran PC. 2016. Interference-driven spacer acquisition is dominant over naive and primed adaptation in a native CRISPR-Cas system. Nat Commun 7:12853. PMC5059440

50. Stern A, Keren L, Wurtzel O, Amitai G, Sorek R. 2010. Self-targeting by CRISPR: gene regulation or autoimmunity? Trends Genet 26:335–40. PMC2910793

51. Cui L, Bikard D. 2016. Consequences of Cas9 cleavage in the chromosome of *Escherichia coli*. Nucleic Acids Res 44:4243–51. PMC4872102

52. Stachler AE, Turgeman-Grott I, Shtifman-Segal E, Allers T, Marchfelder A, Gophna U. 2017. High tolerance to self-targeting of the genome by the endogenous CRISPR-Cas system in an archaeon. Nucleic Acids Res 45:5208–5216. PMC5435918

53. Xu T, Li Y, Shi Z, Hemme CL, Li Y, Zhu Y, Van Nostrand JD, He Z, Zhou J. 2015. Efficient Genome Editing in *Clostridium cellulolyticum* via CRISPR-Cas9 Nickase. Appl Environ Microbiol 81:4423–31. PMC4475897

54. Guan J, Wang W, Sun B. 2017. Chromosomal Targeting by the Type III-A CRISPR-Cas System Can Reshape Genomes in *Staphylococcus aureus*. mSphere 2. PMC5687920

55. Dy RL, Pitman AR, Fineran PC. 2013. Chromosomal targeting by CRISPR-Cas systems can contribute to genome plasticity in bacteria. Mob Genet Elements 3:e26831. PMC3827097

56. Canez C, Selle K, Goh YJ, Barrangou R. 2019. Outcomes and characterization of chromosomal self-targeting by native CRISPR-Cas systems in *Streptococcus thermophilus*. FEMS Microbiol Lett 366.

57. Westra ER, Levin BR. 2020. It is unclear how important CRISPR-Cas systems are for protecting natural populations of bacteria against infections by mobile genetic elements. Proc Natl Acad Sci U S A 117:27777–27785. PMC7668106

58. Rollie C, Chevallereau A, Watson BNJ, Chyou TY, Fradet O, McLeod I, Fineran PC, Brown CM, Gandon S, Westra ER. 2020. Targeting of temperate phages drives loss of type I CRISPR-Cas systems. Nature doi:10.1038/s41586-020-1936-2.

59. van Houte S, Ekroth AK, Broniewski JM, Chabas H, Ashby B, Bondy-Denomy J, Gandon S, Boots M, Paterson S, Buckling A, Westra ER. 2016. The diversity-generating benefits of a prokaryotic adaptive immune system. Nature 532:385–8. PMC4935084

60. Watson BNJ, Steens JA, Staals RHJ, Westra ER, van Houte S. 2021. Coevolution between bacterial CRISPR-Cas systems and their bacteriophages. Cell Host Microbe 29:715–725.

61. Vercoe RB, Chang JT, Dy RL, Taylor C, Gristwood T, Clulow JS, Richter C, Przybilski R, Pitman AR, Fineran PC. 2013. Cytotoxic chromosomal targeting by CRISPR/Cas systems can reshape bacterial genomes and expel or remodel pathogenicity islands. PLoS Genet 9:e1003454. PMC3630108

62. Aklujkar M, Lovley DR. 2010. Interference with histidyl-tRNA synthetase by a CRISPR spacer sequence as a factor in the evolution of *Pelobacter carbinolicus*. BMC Evol Biol 10:230. 2923632

63. Selle K, Klaenhammer TR, Barrangou R. 2015. CRISPR-based screening of genomic island excision events in bacteria. Proc Natl Acad Sci U S A 112:8076–81. PMC4491743

64. Palacios Araya D, Palmer KL, Duerkop BA. 2021. CRISPR-based antimicrobials to obstruct antibiotic-resistant and pathogenic bacteria. PLoS Pathog 17:e1009672. PMC8266055

65. Bae T, Kozlowicz B, Dunny GM. 2002. Two targets in pCF10 DNA for PrgX binding: their role in production of Qa and *prgX* mRNA and in regulation of pheromone-inducible conjugation. J Mol Biol 315:995–1007.

66. Manson JM, Keis S, Smith JMB, Cook GM. 2003. A Clonal Lineage of VanA-Type *Enterococcus faecalis* Predominates in Vancomycin-Resistant Enterococci Isolated in New Zealand. Antimicrobial Agents and Chemotherapy 47:204–210.

67. Gold OG, Jordan HV, van Houte J. 1975. The prevalence of enterococci in the human mouth and their pathogenicity in animal models. Arch Oral Biol 20:473–7.

68. Perez-Casal J, Caparon MG, Scott JR. 1991. Mry, a trans-acting positive regulator of the M protein gene of *Streptococcus pyogenes* with similarity to the receptor proteins of two-component regulatory systems. J Bacteriol 173:2617–24. PMC207828

